# Prevalence of Problematic Papers in Non-Coding RNA Research

**DOI:** 10.1101/2024.08.28.607530

**Authors:** Ying Lou, Zhengyi Zhou, Zhesi Shen, Menghui Li

## Abstract

This study examines the prevalence of problematic papers in the rapidly growing field of non-coding RNA (ncRNA) research. Analysis of 153,826 ncRNA papers during 2000-2023 reveals that around 1.79% have been retracted and an additional 5.68% have raised concerns on PubPeer. The number of problematic papers has steadily increased, peaking in 2019 when the concerning and retraction rates reached nearly 10.8% and 3.7%, respectively. These unreliable papers have been widely disseminated, accumulating hundreds of thousands of citations in academic literature, patents, clinical trials, and policy documents, posing a significant threat to research integrity and public health. The main issues identified include image manipulation, data falsification, fake peer reviews, and ethical lapses. The findings call for urgent, comprehensive scrutiny of ncRNA publications and broader reforms to address systemic problems driving the proliferation of problematic research.

---

*The study underscores the prevalence of problematic non-coding RNA papers, calling for the critical necessity for heightened scrutiny to identify and address these issues.*

Non-coding RNA (ncRNA) plays a crucial role as potent regulators of numerous biological functions in the detection and treatment of various diseases [1]. The study of ncRNA is thriving due to a large amount of under-investigated ncRNA [2], leading to a ‘wild’ growth in publications [3]. For example, the number of publications has risen from 102 in 2000 to 18,491 in 2021 (Figure 1A). Nevertheless, some under-investigated ncRNAs are being actively exploited for potentially fraudulent research [4]. For instance, over 8,700 ncRNA-related papers have been questioned on PubPeer, with more than 2,700 papers retracted, making ncRNA one of the most severely impacted areas by academic misconduct [5]. Furthermore, future research may be misdirected by the problematic papers [4,6]. Even more concerning is that treatments based on problematic conclusions put patients at risk [6]. When many problematic papers emerge within a field, it doubts the credibility of all papers within that field. This research uncovered a significant prevalence of unreliable articles in the ncRNA field, which have been extensively circulated. Therefore, it is crucial and urgent to raise awareness about the severity of problematic papers in the ncRNA field and to advocate for stronger scrutiny of all publications.

**Figure 1:**
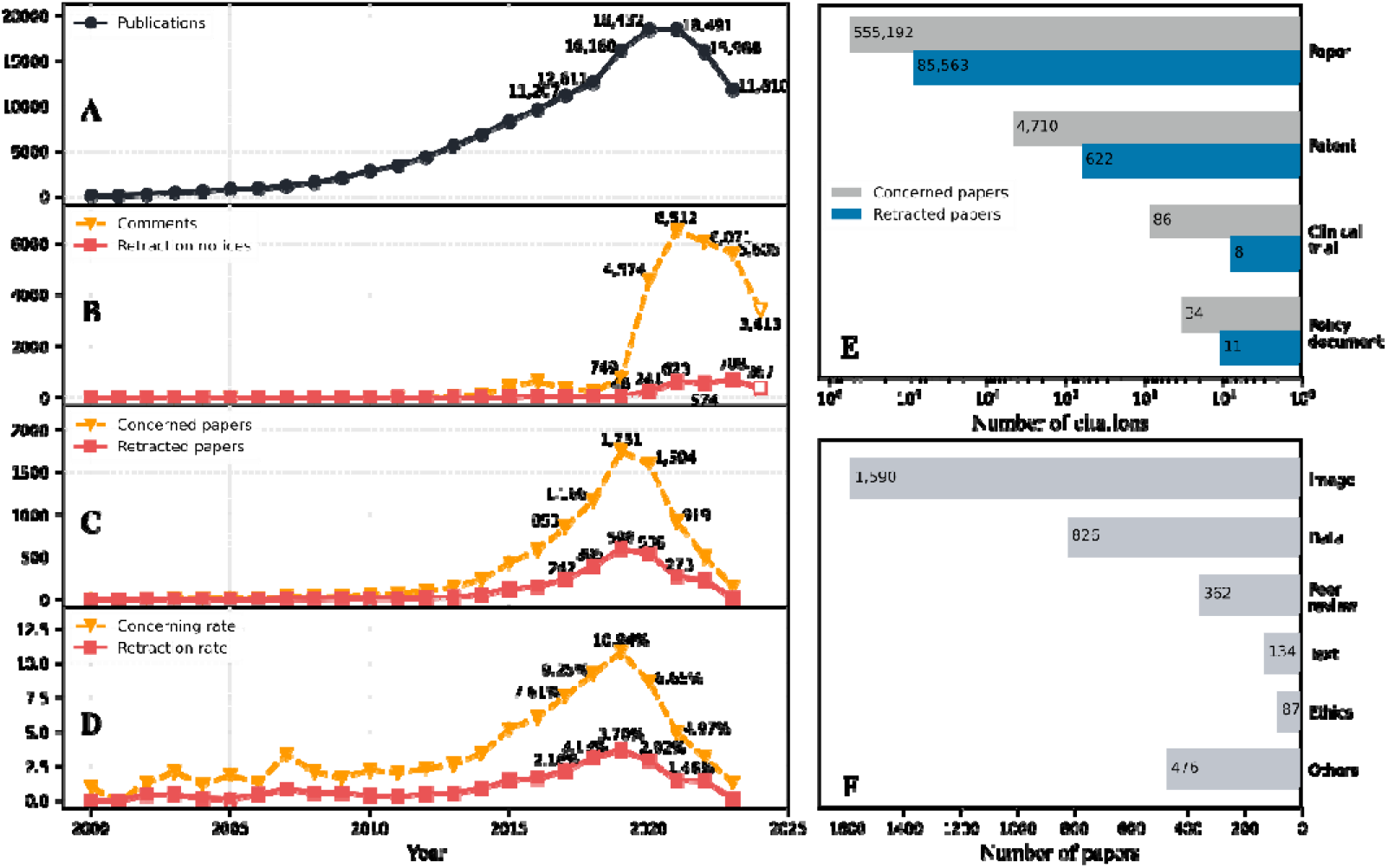
Characterizing the severity of problematic papers in the field of non-coding RNA. (A) The number of papers published each year. (B) The number of comments on PubPeer and retraction notices issued by journals each year. (C) The number of concerned papers and retracted papers for the year of publication. (D) The concerning rate and retraction rate for the year of publication. (E) The wide dissemination, as measured by the number of citations in papers, patents, clinical trials, and policy documents, for both the concerned papers and the retracted papers. (F) The top integrity issues, measured by the number of related retracted papers.

## The proportion of problematic papers is alarmingly high

This analysis covers 153,826 papers on the topics of Micro & Long Noncoding RNA in the InCites database between 2000 and 2023 (Accessed on July 1^st^, 2024), considering only Articles, Reviews, and Retracted Publications. Among these papers, 2,748 papers have been retracted for various reasons, accounting for around 1.79% of the total publications. In addition, 8,739 papers have sparked concerns on PubPeer, constituting approximately 5.68% of the total. These papers have received more than 28,800 comments in PubPeer, almost all questioning their contents [7].

The number of comments on PubPeer surged from 749 in 2019 to 6,512 in 2021, consistently maintaining a high level of over 5,600 comments every year. During the same period, the number of retractions experienced a rapid increase, ultimately peaking at over 700 retractions by 2023 (Figure 1B). With the substantial increase in comments or retractions, it casts a shadow of doubt over the overall credibility of ncRNA papers.

From a historical perspective on publication years, the quantity of concerned papers and retracted papers appears intricately linked to the overall volume of publications. Specifically, the count of concerned or retracted papers ascended from 3 or 1 in 2002 to 1,751 or 598 in 2019, respectively (Figure 1C). Moreover, a consistent upward trend in the annual proportions of both concerned and retracted papers is observed. The peaks observed in 2019, with a concerning rate of 10.84% and a retraction rate of 3.70%, highlight a significant portion of papers being highly problematic (Figure 1D). These findings expose a troubling pattern of increasing proportions of questionable research outputs over the years, highlighting the need for special attention to publications around 2019.

## Unreliable knowledge has been widely disseminated, posing a threat to life and health, highlighting the critical need for urgent scrutiny

The retracted papers have garnered over 85,500 citations in academic papers, more than 600 citations in patents, been cited in clinical trials 8 times, and referenced in policy documents 11 times (Figure 1E). In addition, the papers concerned on PubPeer have accumulated over 555,100 paper citations, more than 4,700 patent citations, over 80 clinical trial citations, and over 30 policy document citations (Figure 1E). For instance, a retracted paper^1^ had amassed more than 2,000 paper citations and had been referenced in patents over 130 times. Additionally, it was cited in 4 clinical trials and mentioned in 3 policy documents in Dimensions (Accessed on August 26th, 2024). Similarly, another paper, concerned on PubPeer in 2019 for irreproducible results^2^, had garnered over 4,500 citations and had been referenced in patents 52 times. These findings suggest that the dissemination of unreliable knowledge has been alarmingly extensive, spanning from basic research to clinical applications and policy formulation, posing a particularly dangerous threat to life and health.

## Scrutiny can start with these characteristics

What characteristics will help us to identify problematic papers? By analyzing and compiling the retraction reasons [8], we find the primary issues are related to image, data, peer review, text, and ethics (Figure 1F). In particular, image-related problems (including image reuse and falsification/fabrication) have emerged as the most common issue which affects 1,590 papers, representing a significant 57.86% of retractions. Meanwhile, within the papers commented on PubPeer, 7,307 are linked to image-related concerns, making up 83.61% of the total cases. Data-related concerns, e.g., data reuse and data falsification/fabrication, have affected 826 papers. Fake peer review, text reuse, and lack of ethical approval are the other three leading issues. Based on the above characteristics, post-publication scrutiny can commence with the help of AI-based tools like ImageTwin, ImaCheck, and Proofig to detect suspicious images [9], as well as STM Integrity Hub to identify Paper Mill [10], and iThenticate to check text similarity.

## Conclusion and discussion

This study examines the prevalence of problematic papers during the flourishing research in the ncRNA field. Approximately 9.2% of papers published during 2017-2020 have been commented on PubPeer, with 3.04% of papers from the same period having been retracted. Moreover, the widespread dissemination of problematic papers not only undermines the research integrity but also misdirects the subsequent studies. The issues revealed in ncRNA are just the tip of the iceberg. Problematic papers have already permeated all fields [5].

It is generally believed that the prevalence of problematic papers is often linked to the pervasive ‘publish or perish’ pressure, highlighting a systematic problem within academia. Maintaining scientific integrity necessitates a pause to address this challenge strategically. All stakeholders bear the responsibility of thoroughly scrutinizing publications. While it is essential to prevent the publishing of new problematic papers, it is equally vital to scrutinize those already published, which can mitigate the influence of unreliable knowledge on both the academic community and the public.

## Data and Methods

About 150,000 papers categorized under the meso-level Citation Topics -- Micro & Long Noncoding RNA (ncRNA) from the InCites database were retrieved on July 1, 2024, covering the publication period from 2000 to 2023. The dataset included articles, reviews, and retracted publications.

To identify problematic papers, we collected the retraction data from the Amend platform[8] and comments on PubPeer. The dissemination of problematic papers was assessed through citation analysis. The citations from papers, patents, policy documents and clinical trials are accessed from Dimensions. Based on the retraction notice recorded in Amend, we compiled retraction reasons and categorized issues into image related, data related, peer review related, text related, and ethical related categories. The full workflow is illustrated in Fig.2. The dateset can be accessed from https://zenodo.org/doi/10.5281/zenodo.13383979.

**Figure 2.**
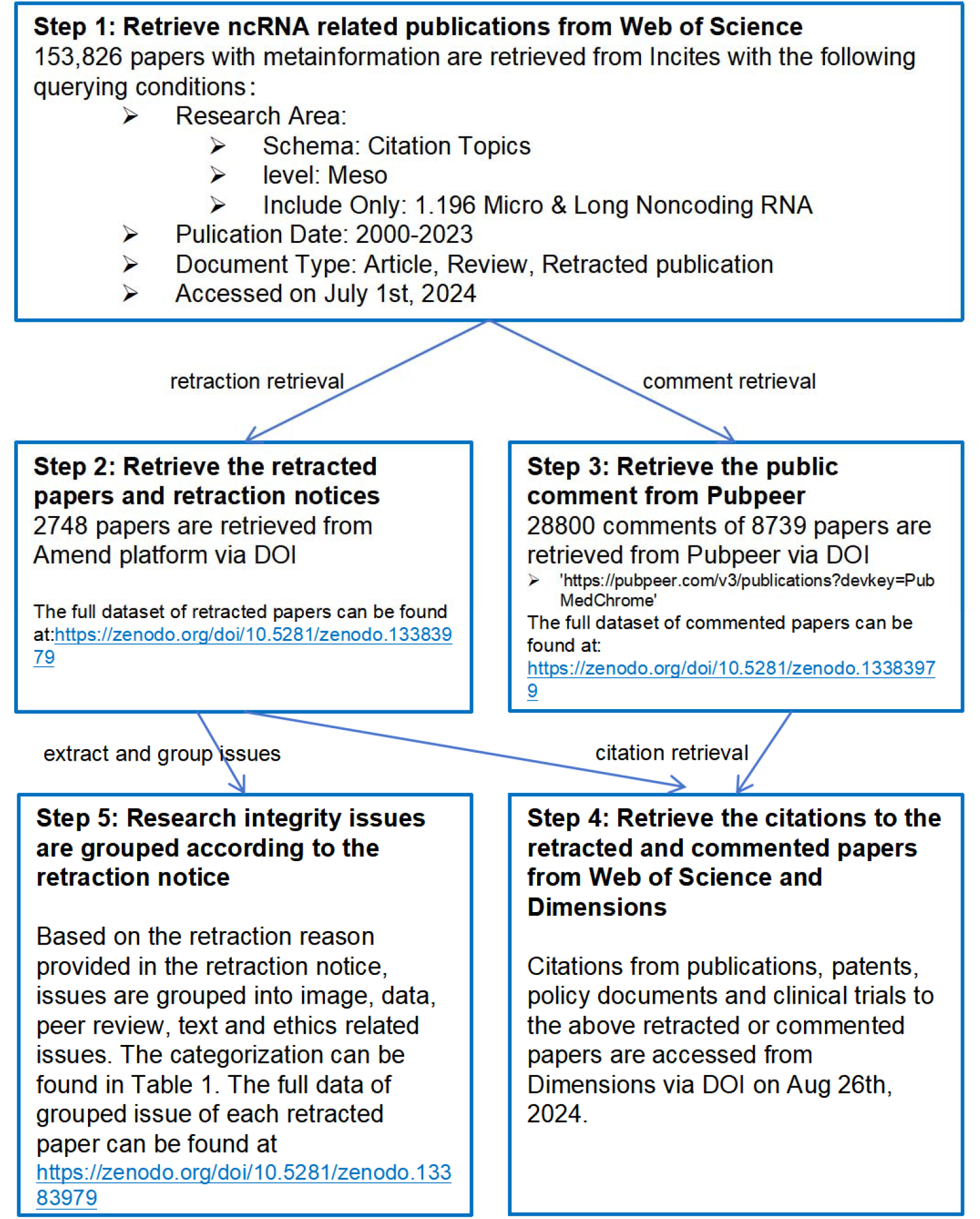
Workflow of data collection and processing.

## Acknowledgments

This study is partially supported by the National Natural Science Foundation of China (grant no. 71974017) and the LIS Outstanding Talents Introducing Program, Bureau of Development and Planning, CAS (2022).

## Statement

During the preparation of this work the author(s) used LLM in order to improve readability and language. After using this tool/service, the author(s) reviewed and edited the content as needed and take(s) full responsibility for the content of the publication.

---

**Table 1.** Grouped issues based on retraction reasons.

| Issue Group | Retraction Reasons |
| --- | --- |
| Image | Image Falsification; Image Plagiarism; Image Fabrication; Image Duplication; Image Error |
| Data | Data Falsification; Data Plagiarism; Data Fabrication; Data Duplication; Data Error |
| Peer Review | Fake Peer Review |
| Text | Text Plagiarism; Text Duplication; Text Error |
| Ethics | Lack of IRB/IACUC Approval; Lack of Informed/Patient Consent; Privacy leak; Lack of trial registration |
| Others | Other reasons except the above reasons |

## Footnotes

1 10.1016/j.ygyno.2008.04.033

2 https://pubpeer.com/publications/9E5F06FC1E57F825881B4FC37956D4

